# HCN channels sense temperature and determine heart rate responses to heat

**DOI:** 10.1101/2023.09.02.556046

**Authors:** Yuejin Wu, Qinchuan Wang, Jonathan Granger, Oscar Reyes Gaido, Eric Nunez Aguilar, Andreas Ludwig, Anna Moroni, Mario A. Bianchet, Mark E. Anderson

## Abstract

Heart rate increases with heat, [1–3] constituting a fundamental physiological relationship in vertebrates. Each normal heartbeat is initiated by an action potential generated in a sinoatrial nodal pacemaker cell. Pacemaker cells are enriched with hyperpolarization activated cyclic nucleotide-gated ion channels (HCN) that deliver cell membrane depolarizing inward current that triggers action potentials. HCN channel current increases due to cAMP binding, a mechanism coupling adrenergic tone to physiological ‘fight or flight’ heart rate acceleration. However, the mechanism(s) for heart rate response to thermal energy is unknown. We used thermodynamical and homology computational modeling, site-directed mutagenesis and mouse models to identify a concise motif on the S4-S5 linker of the cardiac pacemaker HCN4 channels (M407/Y409) that determines HCN4 current (I*_f_*) and cardiac pacemaker cell responses to heat. This motif is required for heat sensing in cardiac pacemaker cells and in isolated hearts. In contrast, the cyclic nucleotide binding domain is not required for heat induced HCN4 current increases. However, a loss of function M407/Y409 motif mutation prevented normal heat and cAMP responses, suggesting that heat sensing machinery is essential for operating the cAMP allosteric pathway and is central to HCN4 modulation. The M407/Y409 motif is conserved across all HCN family members suggesting that HCN channels participate broadly in coupling heat to changes in cell membrane excitability.

## Introduction

Hyperpolarization activated cyclic nucleotide-gated ion channels (HCN1-4) govern cell membrane excitability in neurons, smooth muscle and heart. All HCN family channels are equipped with a cyclic nucleotide binding domain.[4] HCN4, the predominant HCN channel expressed in cardiac sinoatrial nodal (SAN) pacemaker cells, initiates each normal heart beat by a cell membrane depolarizing inward current (I*_f_*)[5] that triggers an action potential. The SAN cell I*_f_* is augmented in response to extracellular catecholamines by an adrenergic receptor mediated increase in cyclic adenosine monophosphate (cAMP), an intracellular second messenger that signals through the HCN4 cyclic nucleotide binding domain. cAMP increases I*_f_* to accelerate SAN action potentials and heart rate, as part of a physiological ‘fight or flight’ response.

However, heart rates are also increased by heat in humans and in mice, constituting a conserved stress response operated by an unknown mechanism.[6] Heart rate increases with temperature with a Q_10_∼2.[2, 7] Understanding the interrelationship between heat, cell membrane excitability and heart rate is an important goal for the life sciences[8], in part, because of rising temperatures worldwide [9]. Based on the importance of HCN4 in heart rate acceleration and the ability of certain ion channels to respond to heat,[10] and to the known temperature dependency of I*_f_* in SAN [11] and Purkinje fibers[12] we asked if HCN4 contributed to heart rate responses to heat.

We identified a heat resistant mutant (M407Q/Y409F) where I*_f_* lacked augmentation by heat. In order to determine if the M407/Y409 site was important for heart rate, we developed a mouse SAN cell model capable of inducible deletion of *Hcn4* and replacement expression of M407Q/Y409F *HCN4* mutant or WT channels. SAN cells with replacement expression of WT human HCN4 exhibited normal rate responses to heat, but SAN cells expressing M407Q/Y409F human HCN4 mutants lacked physiological coupling of heart rate increases to heat. We used CRISPR/Cas 9 to generate mice with M407Q/Y409F mutations in *Hcn4.* Homozygous mice were not viable, but mice with heterozygous expression of *Hcn4* M407Q/Y409F mutants were born in Mendelian ratios. SAN cells and isolated hearts from M407Q/Y409F heterozygous mice had depressed rate responses to heat. Our data show an unanticipated role of HCN4 channels in driving heart rate increases to heat. Because the M407/Y409 motif is conserved across all HCN family ion channels, our findings suggest that HCN channels participate broadly in thermal contributions to cell membrane excitability.

## Results

### Heat-induced rate increases in cardiac pacemaker cells require HCN4 and I_f_

Because the basis for heat induced heart rate increases is unknown, we first asked if spontaneous action potential rates in current clamped isolated SAN cells (Fig 1a) are thermally responsive. We quantified SAN responses to heat using an established temperature coefficient, Q_10_ (Q_10_ = (R2/R1)^10C/(T2-T1)^) where R is the rate of spontaneous action potentials measured at a given temperature, T. In vertebrates the Q_10_ for heart rate in vivo is ∼2.0,[13] equivalent to doubling of heart rate over a 10°C temperature range. We found that spontaneous action potential rates increase with rising temperature in SAN cells isolated from mice (Fig 1b) and have a Q_10_ of 1.97± 0.07, similar to heart rate responses to heat in vivo, in humans and other vertebrates.[7, 13-15]

**Figure 1.**
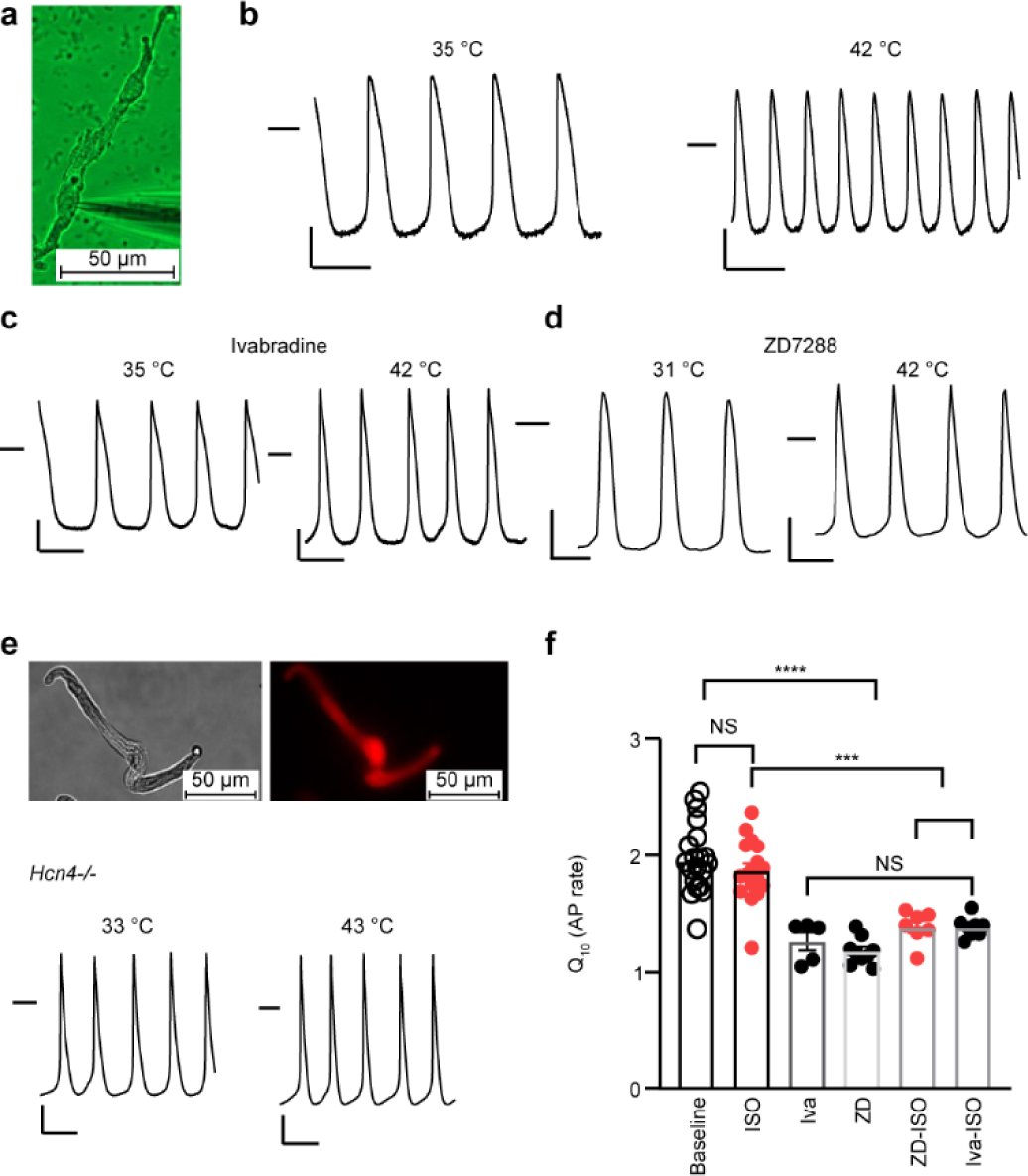
HCN4 antagonists or *Hcn4* knock out prevent action potential increases by heat in SAN cells. **a,** Isolated SAN cell in whole cell mode current clamp configuration, scale bar=50 µm. **b**, Representative SAN cell action potential (AP) tracings recorded under lower (left) and increased temperature (right). **c**, Representative SAN cell AP tracings recorded in the presence of HCN4 antagonist agents Ivabradine (4 μM) or **d**, ZD7288 (4 μM) under lower (left) and increased temperature (right). **e**, upper panel, Example of adenovirus infected SAN cells from floxed mice expressing mCherry and Cre (red, right) (scale bar=50 µm). lower panel, Representative AP tracings recorded under lower basal (left) and increased temperature (right) conditions from cultured single SAN cells isolated from *Hcn4^flox/flox^* mice cultured in the presence of adenovirus expressing Cre recombinase (see Methods). **b-e,** the horizontal lines mark 0 mV. Scale bars are 200 ms horizontal and 20 mV vertical. **f,** Summary data for Q_10_ from SAN cell AP rate responses to temperature increases. The presence of isoproterenol (ISO, 1 μM), Ivabradine (Iva, 4 μM) or ZD7288 (ZD, 4 μM) in the bath solution is indicated. One-way ANOVA, ***p<0.001, ****p<0.0001, n=5-20/group.

Two I*_f_* antagonist drugs (ivabradine [16] and ZD7288 [17]) prevented SAN thermal rate responses (Q_10_ = 1.27±0.08 for ivabradine and 1.18±0.04 for ZD7822, p<0.001 as compared to control condition, Fig 1c and d), without affecting rate increases from isoproterenol (Fig S1a), a β-adrenergic receptor agonist that accelerates heart rate by HCN4 and cAMP dependent and independent pathways.[18] We cultured SAN cells isolated from mice where *Hcn4* was flanked by LoxP insertions (*Hcn4*^L2/L2^).[19] These SAN cells were infected with adenovirus expressing Cre recombinase and a red fluorescent protein marker (mCherry, see Methods, Fig 1e) for 24 hours to generate *Hcn4* knock out (*Hcn4^-/-^*) SAN cells. These *Hcn4*^-/-^ SAN cells lacked HCN4 current, I*_f_* (Fig S1b), indicating successful excision of *Hcn4* by Cre recombinase. Consistent with our findings using I*_f_* antagonist drugs, cultured *Hcn4^-/-^* SAN cells lacked thermal responses (Q_10_ = 1.23±0.06, Fig 1e).

In addition to I*_f_*, Calmodulin kinase II (CaMKII) activation[18] contributes to SAN cell rate increases and is potentially sensitive to temperature. SAN cells with CaMKII inhibition due to transgenic expression of an inhibitory peptide, AC3-I, have diminished rate responses to isoproterenol.[18] However, we found that AC3-I expressing SAN cells showed similar rate increases in response to temperature compared to SAN cells isolated from WT littermates (Fig S1c). Taken together (Fig 1f), these findings supported a concept that SAN cell action potential rates were increased by thermal energy, similar to in vivo heart rate responses, and suggested that I*_f_* was responsible for all or most of SAN cell heat sensitivity.

### I_f_ has a Q_10_ near 2.0 that is independent of cAMP signaling

Given our findings showing HCN4 and I*_f_* were essential for heat mediated SAN cell action potential frequency increases, we next performed voltage clamp experiments on isolated SAN cells and found I*_f_* was enhanced by increasing temperatures between 30°C-45°C with a Q_10_ (Q_10_ is calculated based on current density change by temperature) of around 2 (e.g. 2.15±0.11 at −60 mV, 1.94±0.30 at −130 mV) at cell membrane test potentials from −60 mV to −130 mV (Fig 2a and Fig S1d). In order to determine if the I*_f_* heat response was related to SAN cell specific elements, we transfected (human) HCN4 cDNA into human embryonic kidney 293 (HEK) cells. The HCN4 channel current recorded from HEK cells increased with heat with a Q_10_ of around 2 (e.g. 2.12±0.18 at −60 mV, 2.02±0.22 at −130 mV) at test potentials between −60 mV and −130 mV, similar to our measurements in SAN cells. (Fig 2b and Fig S1d). We chose to focus on Q_10_ values at −60 mV, a physiologically relevant cell membrane potential for SAN cells. The Q_10_ for I*_f_* recorded from HEK cells was similar in the presence or absence of cAMP dialysis (Fig 2c) while ivabradine and ZD7288 prevented the I*_f_* response to heat in SAN cells with or without ISO (Fig 2d). Similar to SAN action potential rates, heat increased I*_f_*, apparently by a mechanism that is independent of cAMP sensing.

**Figure 2.**
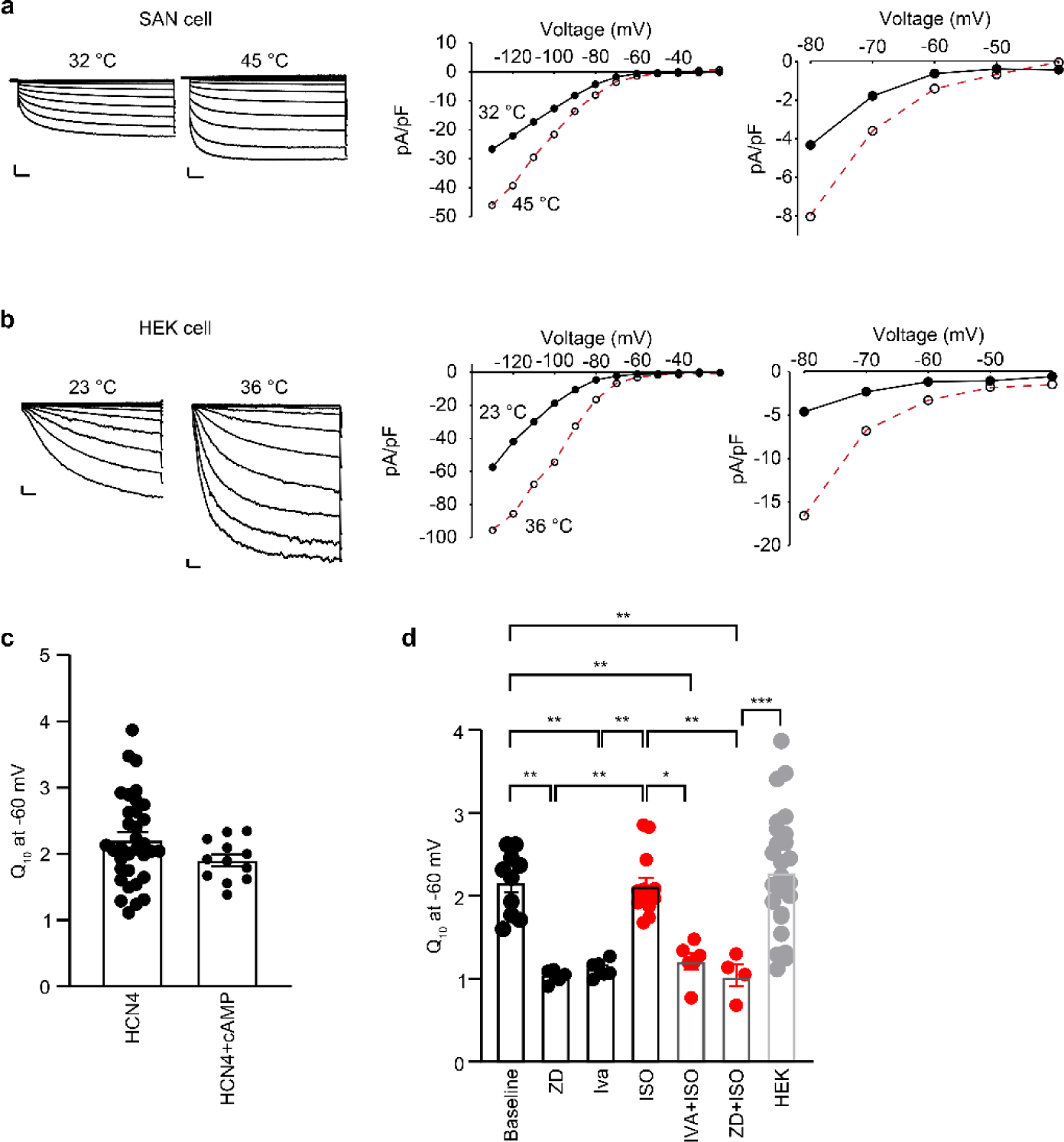
HCN4 current (I*_f_*) has a Q_10_ of ∼2.0 that is independent of cAMP signaling. Representative I*_f_* (HCN4 current) tracings recorded under lower temperature and increased temperature (left panels) conditions from **a,** isolated SAN cells or **b,** in human embryonic kidney cells (HEK) transfected with *HCN4* (see Methods). I*_f_* current-voltage relationship (right panels). The right-most panels are expanded to highlight the physiological cell membrane potential range for SAN cells (−40 to - 80 mV). The horizontal scale bar is 500 ms and the vertical scale bar is 5 pA/pF in both (**a**) and (**b**). **c**, Summary data for Q10 from HCN4 channel currents at −60 mV from HEK cells expressing *HCN4* in the presence or absence of cAMP (n=12-34/group). **d**, Summary Q10 data for I*_f_* at −60 mV from isolated SAN cells treated with ISO (1µM), and/or Ivabradine (Iva, 4 μM) or ZD7288 (ZD, 4 μM) (n=4-13/group). one way ANOVA *p<0.05, **p<0.01, ***p<0.001.

### HCN4 temperature responses rely on a concise intracellular linker motif

Based on these findings showing that HCN4 conducts a heat-responsive I*_f_* we next sought to identify residues modulating the channel thermal responses. The heat dependence of channel activation is a function of the curvature of free energy (ΔΔ*G*(*T*))[20] Such curvature depends on the variation of the specific heat at the pressure constant ΔΔ*C_p_* (*T*) (see Supplementary material and equations 1-6). Δ*C_p_* (*T*) in proteins is mostly a function of the area exposed to the solvent[20] and its variation during channel activation is related to the change in the solvent-accessible surface area (SASA).

To identify candidate amino acids making significant contributions to the HCN4 temperature response, we used a common-sense approach: focusing our search on residues of protein regions exhibiting significative changes in SASA measured between the two states represented by the structures of the cAMP-free (closed) and cAMP-bound (activated) states of the HCN4 channel.

We performed a comparison of the human HCN4 structures with homology models of similar channels, including from Tardigrade (*Ramazzottius varieornatus*), an organism resistant to extreme heat. This analysis helped us to pinpoint by thermodynamic analysis specific residues that modulate HCN4 thermal responses. The HCN4 protein forms a pore as an aggregation of four subunits. Each of the subunits consists of six membrane spanning domains (transmembrane S1-S6) connected by intra- and extracellular linkers. There is a prominent intracellular C terminus that contains the cyclic nucleotide binding domain (Fig 3a) connected to the S6 via a C-linker that forms the “disk domain” (Figure 3a). Binding of cAMP to the cyclic nucleotide binding domain induces conformational changes that are transmitted to the transmembrane region via the C-linker and facilitate pore opening. Even though HCN4 channels require hyperpolarizing voltages to move the voltage sensor and to open the pore, the conformational changes induced by cAMP binding in the cytosolic domain are present in the cryoEM structures of the channel solved in the absence of voltage (0 mV). We define this structure as “activated” in our analysis below. The overlap of the human HCN4 structures, apo/closed (PDB:6GYN) and holo/activated (6GYO), Figure 3b, shows that, relative to the other domains, the transmembrane portion of the channel displays significantly less changes. For example, in the human channel the rmsd differences between transmembrane domains (a.a. 208-530) of apo/closed channel shows a rmsd of 1.05 Å^2^ for backbone atoms, while the disk (a.a. 521-573) has an rmsd of 3.55 Å^2^ and the cyclic nucleotide binding domain (a.a. 574-718) exhibits an rmsd of 3.94 Å^2^. When the same analysis was repeated by comparing the holo structures of human (6GYO) and rabbit (7NP4) HCN4 structures, minor differences emerged in the S4-5 linker that could affect heat sensing. However, we found that HCN4 channel current heat responses were similar between human (Fig 2b and c) and rabbit HCN4 expressed in HEK cells (Fig S2a, and Supplemental Materials). Based on this, we focused on human HCN4 for structural and functional studies.

**Figure 3.**
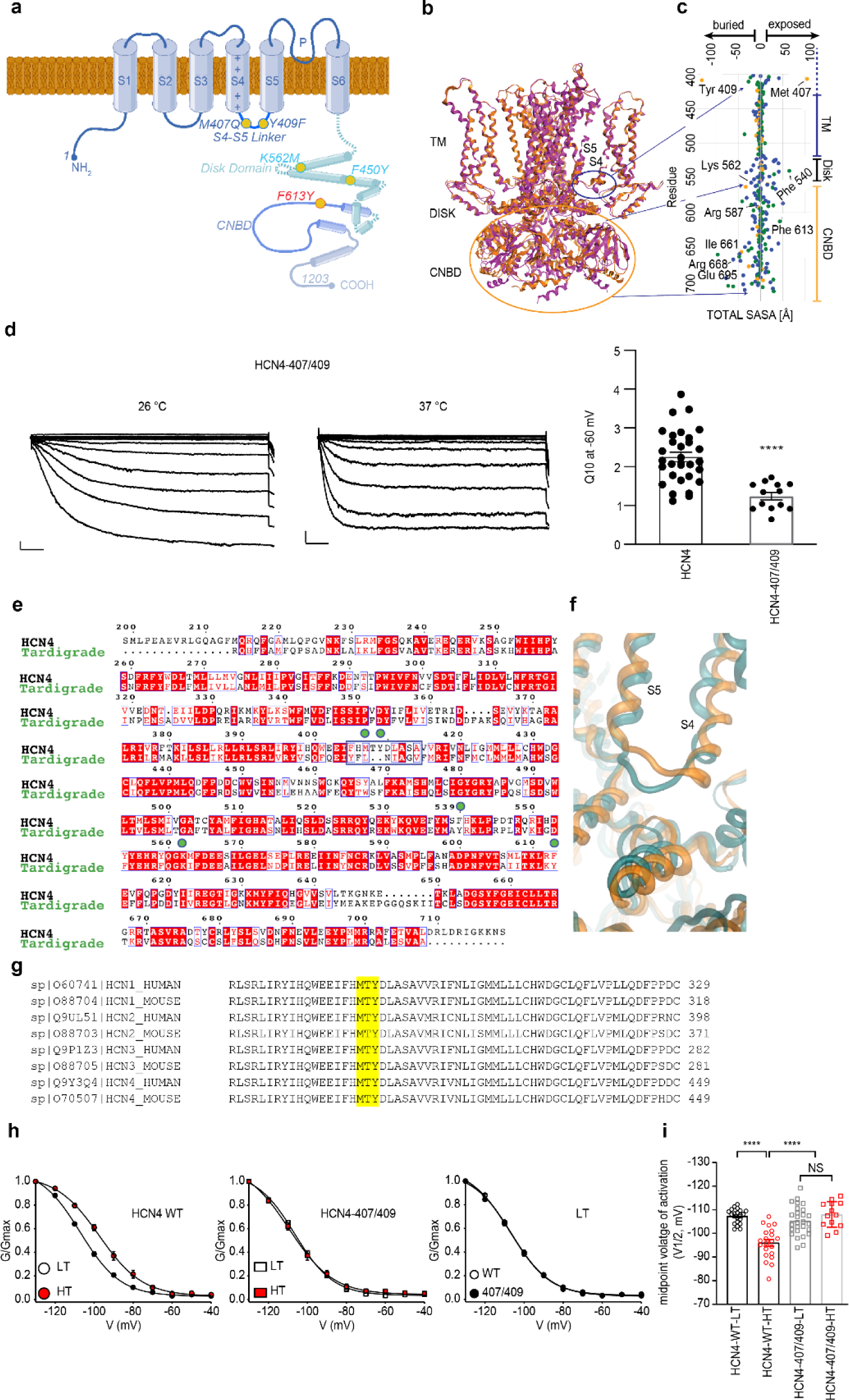
Two amino acids on the 4-5 intracellular linker are essential for heat sensing in HCN4. **a**, Schematic depiction of the subunit structure of HCN4. The pore, voltage-sensing, intracellular linker, disc and cyclic nucleotide binding domains are highlighted. **b,** Ribbon diagram showing superimposed HCN4 in apo/closed and holo/activated (cAMP bound) conformations: 6GYN (apo/closed, in purple) and 6GYO (holo/activated, in orange). **c**, Total solvent accessible surface area (SASA) exposed (right side) or buried (left side) in the residues of the S4-5 intracellular linker (top), disk domain (Disk, middle) and CNBD (cyclic nucleotide binding domain, lower) in response to channel opening, apolar (green), polar (blue) and amphipathic (orange) residues are shown as dots. TM indicates transmembrane domain. **d**, Representative whole cell mode HCN4 currents (left) recorded at lower (left, LT: 24-30 °C) and higher temperature (middle, HT: 34-43°C) from cultured HEK 293 cells expressing *HCN4* heat insensitive mutants (407/409); summary Q_10_ data (right) for HCN4 current recorded from HEK 293 cells expressing wild type and 407/409 *HCN4* (see Methods). Scale bars are 500 ms horizontal and 5 pA/pF vertical, ****p<0.0001, n=13-31/group. **e,** Sequence alignments between human HCN4 and a homologous channel from *Tardigrades*. The candidate temperature sensing residues are marked with rectangles and the blue box indicates the position of the intracellular 4-5 linker. The green dots show the residues targeted for mutation in this study. Amino acid sequence entries are from UniProt databank. **f,** Ribbon diagram comparing the closed state of human HCN4 (gold) and the homologous *Tardigrade* channel (green). **g**, Amino acid sequence alignments comparing the 4-5 intracellular linker for human and mouse HCN1-4. **h,** comparison of HCN4 current mean activation responses to temperature increases from HEK 293 cells expressing *HCN4* (left) or *HCN4*^407/409^ (middle). WT (left) or *HCN4*^407/409^ (middle) HCN4 current activation voltage dependence during baseline (empty symbols) and after increased temperature (filled red symbols). Solid lines show Boltzmann fitting (see Methods). The right panel shows comparison of the voltage dependent of activation for WT and *HCN4*^407/409^ mutants at lower baseline temperature. Solid lines show Boltzmann fitting to the data (see Methods). Values are reported in Table S2. **i**, Summary data for midpoint voltage of activation (V1/2). ****p<0.0001, n=20-21/group. Values are reported in Table S2.

Figure 3c shows that the average change in SASA (<ΔSASA>) of the whole human HCN4 channel during channel activation by cAMP. The average value is 1.3 Å^2^ with a standard deviation (σ(SASA)) of 16.0 Å^2^. We analyzed residues showing absolute values of ΔSASA greater than 3 times σ(SASA) (> 48 Å^2^) between apo/closed and holo-activated states of the human channel (Table S1). The most significant increases in the apolar area exposed during its activation are Met 407 (104 Å^2^), Phe 540 (54.9 Å^2^), and Phe 613 (44.4 Å^2^), and the most significant decreases, due to polar area sequestration, are Tyr 409 (−141.6 Å^2^), Arg 668 (−57.8 Å^2^), and Glu 695 (−77.8 Å^2^).

Met 407 and Tyr 409 are part of the S4-5 intracellular linker that exhibits a large conformational change after activation (Fig 3b and c). We excluded residues in disordered regions (e.g., at the C-terminus of the cyclic nucleotide binding domain) from the analysis, focusing on those observed in the structures. To avoid making mutations affecting the activation mechanism, we did not consider residues coordinating intracellular Mg^2+^ (His 406, Asp 410, Glu 556 and His 552, Fig S2b)[21], nor those participating in cAMP binding, such as Ile 661 or Arg 587, and ignored amino acids forming salt bridges stabilizing one particular state of the channel. The above considerations led to a final list of five candidates with the potential to modify the temperature dependence of I*_f_*: Met 407, Tyr 409, Phe 540, Lys 562, and Phe 613 (see Supplementary Materials). Based on our structural analysis and modelling, and the predicted energy contributions due to its solvation or lack thereof between the closed and cAMP-active HCN4, we designed a set of mutations M407Q, Y409F, F540Y, K562M, and F613Y (see Supplementary Materials), predicted to reduce the activation energy barrier or produce positive contributions to the overall 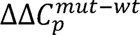, thereby reducing its temperature dependence (lowering Q_10_^mut^/Q_10_^WT^, equation 5, see Supplementary Materials).

Consistent with our predictions, HCN4 channel current recorded from the 5x mutant HCN4 channel was not responsive to temperature between 25-40°C (Fig S2c). We next performed a series of studies to evaluate the thermal responsiveness of HCN4 mutants in isolation and in various combinations (Fig S2d). These studies revealed M407/Y409 as essential and sufficient for conferring thermal responsiveness to HCN4 current (Fig 3d). Based on our finding that thermal sensing in HCN4 substantially resided in the S4-5 intracellular linker region, we asked if this region was important in known temperature sensitive ion channels. Adaptation to temperature in proteins of thermophilic organisms (less sensitive to temperature changes) relies on smaller residue volume, higher hydrophobicity and more charged amino acids, along with a decrease in uncharged polar residues. The Tardigrade hypothetical protein RvY_01629 (GAU89031.1) is a homologous ion channel to HCN4 (54.9% identity). Tardigrada is a phylum of organisms capable of thriving in a wide range of temperatures. Notably, comparison of the HCN homolog from Tardigrade with human HCN4 revealed a two-residue shorter linker between the S4 and S5 helices in Tardigrade (Fig 3e and f). These ‘missing’ amino acids in Tardigrade HCN contribute to a dramatic conformational change during activation in human HCN4 (Supp. Movie 1). The shorter S4-5 linker in Tardigrade HCN suggests a structural basis for diminished heat sensitivity compared to human and mouse HCN4. We interpreted the foreshortened S4-5 linker in the Tardigrade HCN homolog to support our findings that this region is important for heat sensing.

Intriguingly, a region of the S4-5 intracellular linker domain, containing the M407/Y409 residues, was recently identified as important for establishing the melting point for HCN4.[21] This region includes H406 and D410 that are involved in complexing Mg^2+^together with residues from the C linker domain. These contacts are not present in HCN1(5U6P), and are important in transmitting the cAMP effect in HCN4.[21] Because of the potential role of the S4-5 linker in modulating I*_f_* through Mg^2+^ binding, we dialyzed HEK cells expressing HCN4 with EDTA (10 mM) to chelate Mg^2+^, but failed to detect a change in the temperature response (Fig S2e), supporting our modeling predictions that the role of the S4-5 linker in thermal sensing was independent of its role in Mg^2+^ binding. Importantly, M407/Y409 homologous residues, present in the S4-5 intracellular linker domain, are conserved between mouse and humans and are present on all HCN family members (Fig 3g). Currents recorded from HEK cells expressing HCN1 (Q_10_ = 2.25±0.22) and HCN2 (Q_10_ = 2.01±0.14) showed similar thermal responses compared to HCN4 (Fig S3a). Consistent with previous studies, [22, 23] we were unable to efficiently express HCN3 in HEK cell. Mutation of M407/Y409 homologs in human HCN1 (M287/Y289, Q_10_ = 1.10±0.04) and HCN2 (M356/Y358, Q_10_ = 1.02±0.05) eliminated the thermal responses (Fig S3a), demonstrating the general nature of this thermal sensing motif in HCN family channels. We found that temperature increases shifted the half maximal activation voltage (V_1/2_, mV) in a depolarized direction (11.3±1.6 mV), for WT HCN4 expressed in HEK 293 cells (Fig 3h and Table S2) and this action was prevented in the temperature insensitive HCN4 M407Q/Y409F mutants (Fig 3i and Table S2). Notably, the double mutant showed the same V_1/2_ of the WT at low temperature (Fig 3h-i and Table S2), indicating that the mutations do not affect the voltage-gating mechanism of the channel. We found that homologous rabbit HCN4 mutants (M408Q/Y410F) were similarly unresponsive to heat compared to WT (Fig S3b, Fig S2a and Fig 3d). Taken together, these computational and electrophysiological findings showed M407/Y409, a component of the intracellular S4-S5 HCN4 linker, is sufficient to account for I*_f_* increases in response to heat.

### Defective heat-heart rate coupling in M407Q/Y409F Hcn4 knockin mice

We asked if M407/Y409 contributed to heart rate responses to heat. We used CRISPR/Cas9 gene editing to engineer mice with *Hcn4* harboring the M407Q/Y409F mutations (Fig S4 and Supplemental Materials). No homozygous knockin mice were born from *Hcn4*^+/QF^ x *Hcn4*^+/QF^ crosses (10 *Hcn4*^+/+^, 18 *Hcn4*^+/QF^, and 0 *Hcn4^QF/QF^* pups), suggesting that HCN4 containing WT M407/Y409 is required for development. However, heterozygous *Hcn4* mutant mice (*Hcn4*^+/QF^) were viable and born in ∼1:1 proportion to WT littermates from *Hcn4*^+/QF^ x *Hcn4*^+/+^ crosses (113 *Hcn4^+/+^* and 116 *Hcn4*^+*/QF*^ pups). The I*_f_* in SAN cells isolated from *Hcn4*^+/QF^ mice showed reduced responses to heat compared to I*_f_* recorded from WT littermate control SAN cells (Fig 4a). Similar to I*_f_*, action potential rates recorded from *Hcn4^+/QF^* SAN cells (see Methods) had reduced Q_10_ (Fig 4b). We next recorded electrocardiograms from isolated, coronary artery perfused hearts from WT and *Hcn4*^+/QF^ mice in order to measure heart rate (Fig 4c). The *Hcn4*^+/QF^ heart rates had lower Q_10_ than hearts from WT littermate controls (*Hcn4^+/+^*). Taken together, studies in *Hcn4*^+/QF^ mice supported a role for HCN4 coupling heat to heart rate in SAN cells and ex vivo hearts.

**Figure 4.**
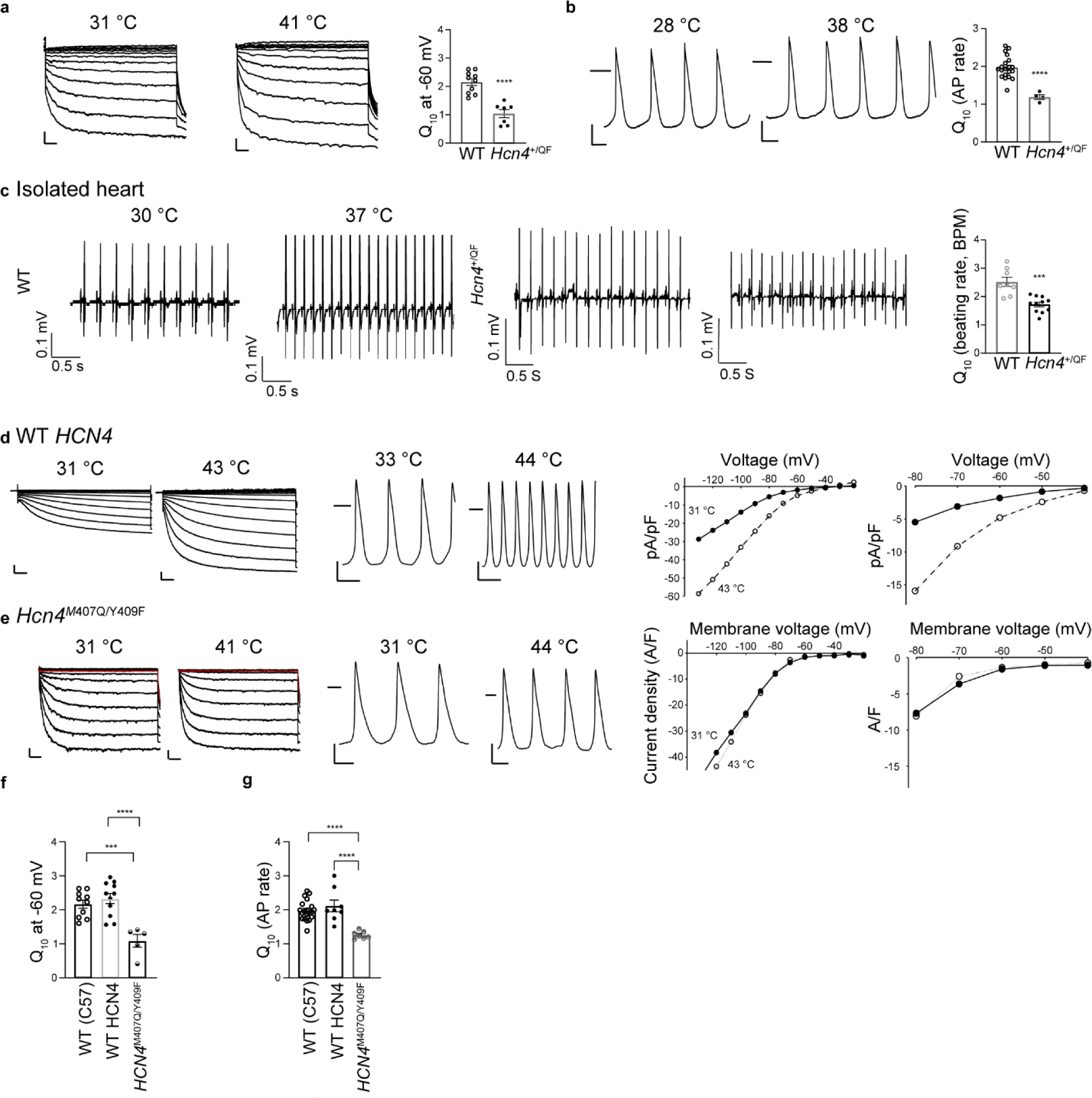
Mice with Hcn4^407/409^ mutant channels lack physiological responses to heat. **a,** Example of I*_f_* current tracings recorded under lower temperature (left) and increased temperature (middle) conditions from isolated single SAN cells of *Hcn4*^+/QF^ mice. Right bar graph is the summary of Q_10_ calculations based on current density at −60 mV from *Hcn4*^+/QF^ mice and WT mice. **b,** Example of action potentials (AP) recorded under lower temperature (left) and increased temperature (middle) conditions from isolated single SAN cells of *Hcn4*^+/QF^ mice. Right bar graph is the summary of Q_10_ calculations based on action potential rates from *Hcn4*^+/QF^ mice and WT mice. **c,** Example of ECG tracings recorded under lower temperature (left) and increased temperature (middle) conditions from isolated hearts of littermate WT control (upper) or *Hcn4*^+/QF^ (lower) mice. Right bar graph is the summary of Q_10_ calculations based on ECG rates (beats/min, BPM) from *Hcn4*^+/QF^ mice and littermate control WT mice. **d,** Upper panels show examples of I*_f_* currents recorded under baseline (left) and increased temperature (middle) conditions from cultured *Hcn4^flox/flox^* SAN cells infected with Ad-mCherry-Cre and Ad-*HCN4*-WT. Scale bar is 5 pA/pF (vertical) and 500 ms (horizontal). I*_f_* current density and voltage relationship between 0 to −140 mV (right, middle) and between −40 to −80 mV (right). All I*_f_* current examples are from the same cell. Full symbols = 31°C, open symbols = 43 °C Lower panels show example AP recordings under basal (left) and increased temperature (right) conditions from cultured SAN cells isolated from *Hcn4^flox/flox^* mice infected with Ad-Cre-*HCN4*-WT. The horizontal line marks 0 mV. Scale bars are 200 ms (horizontal) and 20 mV (vertical). **e,** Upper panels show examples of I*_f_* currents recorded under lower baseline (left) and increased temperature (middle) conditions from cultured *Hcn4^flox/flox^* SAN cells infected with Ad-mCherry-Cre and Ad-*HCN4^QF/QF^*. Scale bar is 5 pA/pF (vertical) and 500 ms (horizontal). I*_f_* current density and voltage relationship between 0 and −140 mV (right, middle) and between −40 to −80 mV (right). I*_f_* recordings and current voltage relationships are from the same cell. Full symbols = 31 °C, open symbols = 41°C Lower panels show example AP tracings recorded under lower basal (left) and increased temperature (right) conditions from cultured single SAN cells isolated from *Hcn4^flox/flox^* mice and infected with Ad-mCherry-Cre and *HCN4^QF/QF^*. The horizontal line marks 0 mV. Scale bars are 200 ms (horizontal) and 20 mV (vertical). **f,** Summary Q_10_ data for I*_f_* currents (n=5-11/group) at −60 mV from cultured *Hcn4^flox/flox^* SAN cells expressing *HCN4* WT, or *HCN4^QF/QF^*. ***p<0.001. **g,** Summary Q_10_ data for AP rates (n=7-8/group) from cultured *Hcn4^flox/flox^* SAN cells expressing *HCN4* WT, or *HCN4^QF/QF^*. ***p<0.001.

We next turned to *Hcn4^L2/L2^* SAN cells[19] infected with Cre expressing adenovirus to develop an exchange system capable of rescuing I*_f_* in an *Hcn4^-/-^* background using cDNAs encoding WT human HCN4 or HCN4^QF^ (see Methods). We performed voltage and current clamp studies using cultured isolated *Hcn4^-/-^* SAN cells expressing WT HCN4 or HCN4^QF^. The SAN cells rescued with WT HCN4 regained I*_f_* and action potential rate responsiveness to heat (Fig 4d), similar to WT SAN cells (Figs 1b and 2a). In contrast, the SAN cells rescued with thermally insensitive HCN4^QF^ mutant channels failed to increase I*_f_* or action potential rate (Fig 4e) in response to heat. The Q_10_ for I*_f_* (Fig 4f) and SAN action potential rate (Fig 4g) after rescue with WT *HCN4* were ∼2.0. Taken together, these data from CRISPR/Cas 9 gene edited mice and SAN cells with *Hcn4* excision and rescue with WT HCN4 or HCN4^Q*F*^ channels showed that M407/Y409 were required for heat responses for SAN action potentials and I*_f_*.

### HCN4 thermal sensing is independent of the cyclic nucleotide binding domain

HCN4 belongs to a family of ion channel proteins that are endowed with and named for a cyclic nucleotide binding domain.[24] Heart rate accelerates as part of the ‘fight or flight’ physiological response due to increased intracellular cAMP that activates HCN4 through the cyclic nucleotide binding domain, causing increased I*_f_* and accelerated action potential initiation.[25] Indeed, SAN I*_f_* increased upon exposure to isoproterenol, a β adrenergic receptor agonist [18, 26], as did the current recorded from HEK cells expressing HCN4 and dialyzed with cAMP (Fig 5a, left panels). Dialysis of cAMP into HEK cells shifted the half maximal activation of the HCN4 current to more depolarized cell membrane potentials (Fig 5a, right panels), an effect similar to that of heat-induced activation of HCN4 (Figure 3I). Furthermore, cAMP and heat both hastened the activation kinetics (Figs 2 and 5), suggesting that these signals may operate on HCN4 to increase the current through a shared mechanism. Based on this observation, we next generated a validated loss of function mutation (EA mutant, containing R669E and T670A) in the *HCN4* cyclic nucleotide binding domain.[27] The EA mutant HCN4 channels failed to increase I*_f_* (Fig 5b, left panels) or affect the half maximal activation voltage (Fig 5b, right panels) upon cAMP exposure. However, EA mutant channels were responsive to increases in temperature, showing elevated peak current and faster activation. Q_10_ was not different from wildtype channels (Fig 5c). Based on these findings, we next asked if cAMP responses required the M407/Y409 motif; I*_f_* recorded from M407Q/Y409F mutants was unresponsive to cAMP (Fig 5d, Table S2). Collectively, we interpret these findings to show that HCN4 ion channels require M407/Y409 to respond to heat and cAMP.

**Figure 5.**
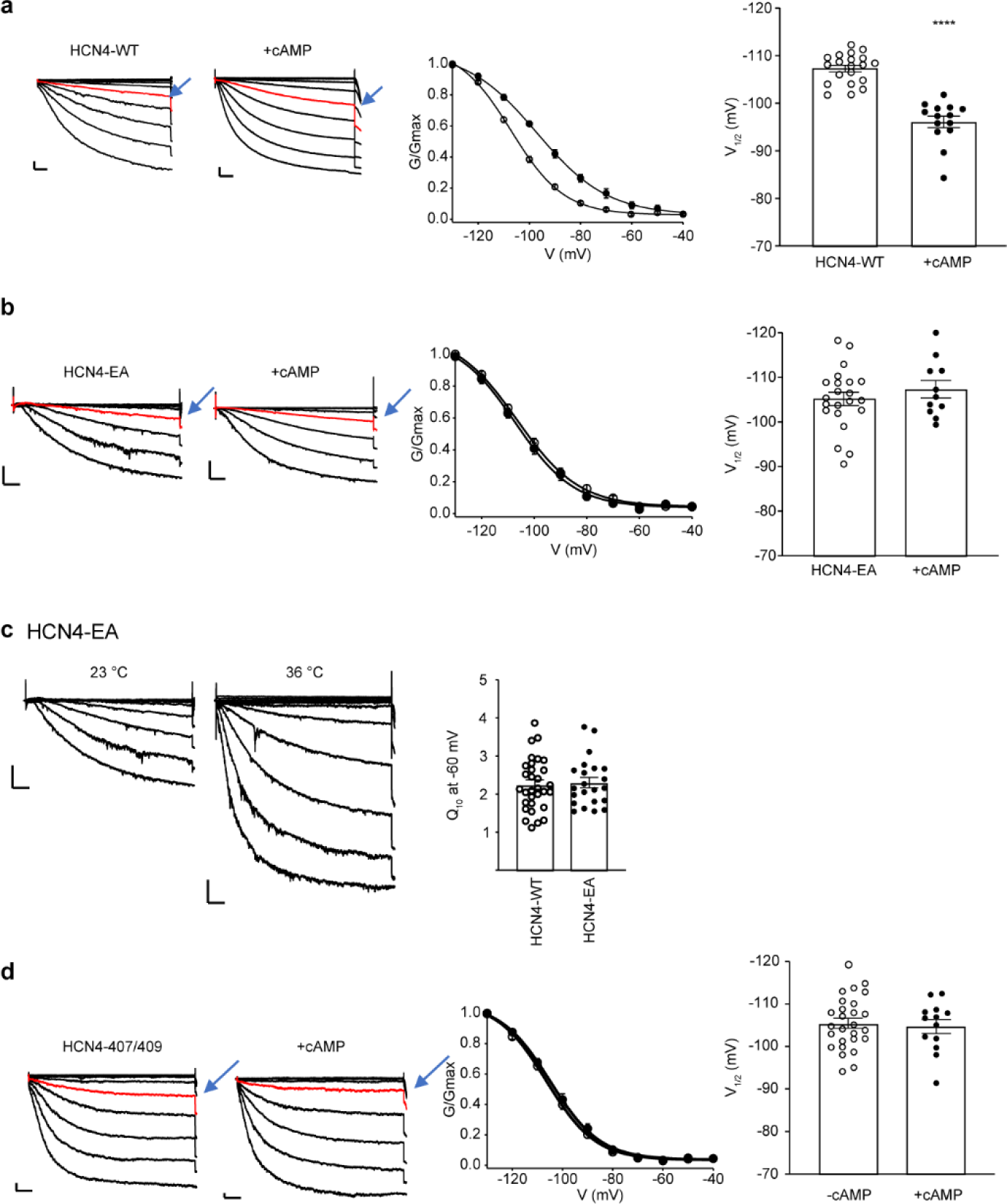
Heat sensing is required for cAMP-dependent activation of HCN4. **a**, Representative I*_f_* currents recorded from *HCN4* transfected HEK 293 cells with or without intracellular dialysis of cAMP (1 mM) (left panel). Boltzmann distribution for *HCN4* transfected HEK 293 cells (middle panel) and summary data for midpoint voltage of activation (right panel, V_1/2_). ****p<0.0001, n=14-20/group. **b,** Representative I*_f_* currents recorded from mutant *HCN4* (EA) transfected HEK 293 cells with or without intracellular dialysis of cAMP (1 mM) (left panel). Data are analyzed and displayed in **b** middle and right panels as in **a** panel (above), n=11-22/group. **c**, Representative I*_f_* currents recorded under lower (left) and increased (middle) temperature conditions from cultured HEK 293 cells expressing *HCN4* EA mutants. Summary Q_10_ data for I*_f_* currents at −60 mV from HEK 293 cells (right panel). ****p<0.0001, n=9-31/group. **d**, Representative I*_f_* currents in the presence or absence of intracellular dialysis with cAMP (1 mM) (left panels) in *HCN4*^407/409^ mutant expressing HEK 293 cells. Right two panels, *HCN4*^407/409^ current activation curves without (empty symbols) and with cAMP (1 mM) (filled symbols) dialysis. Summary data for midpoint voltage of activation (V1/2) with or without cAMP (right), n=13-27/group. Solid lines show Boltzmann fitting to the data (middle, see Methods). Values are reported in Table S2. **a-d,** Scale bars are 500 ms horizontal and 5 pA/pF vertical.

## Discussion

The HCN family of ion channels are defined by a characteristic voltage activated inward current and a cyclic nucleotide response element. Our most complete understanding of the impact of the cAMP response amongst HCN family channels involves the role of HCN4 in heart rate acceleration, a cornerstone of the fight or flight stress response. Up until now, cAMP responses were thought to be the sole mechanism for augmenting HCN channel currents, including the canonical case of I*_f_* underpinning heart rate acceleration. Our findings add a new core concept in cardiac physiology by showing in mice that HCN4 is also responsible for heat triggered increases in heart rate, a feature of all vertebrates.[28] The finding that HCN4 heat sensing relies on two key amino acids of one intracellular linker domain is in line with a recent evidence that Shaker-IR potassium channel also depend upon a homologous intracellular linker domain for thermal responses.[29]. This raises the question whether the (4-5) linker contributes to a general mechanism for heat sensing in ion channels. Although our study has focused on the role of HCN4 in accelerating heart rate, we found that all HCN family members share the critical M407/Y409 residues, albeit with differences in numbering. We found that HCN1 and 2, in addition to HCN4, use the same, concise heat sensing mechanism for augmenting ionic currents. HCN1 and 2 are widely expressed in brain [30], including in regions known to participate in thermal sensing and response. Thus, our work supports future studies to determine if HCN channels contribute more broadly to the systems biology of heat.

Our studies present new evidence that heat sensing is a dominant, but previously unanticipated, mechanism central to the regulation of HCN channels. Loss of M407/Y409 residues, in M407Q/Y409F mutants, prevented heat and cAMP responses. In contrast, a well described loss of function mutation in the cyclic nucleotide binding domain (EA)[27] disabled cAMP responses without affecting heat triggered I*_f_* enhancement or action potential acceleration. Thus, cAMP responses are dispensable to heat responses while the M407/Y409 heat response motif is required for both heat sensing and cAMP responses. In homeothermic organisms core body temperature increases with exercise [31, 32] and fever, [33] both conditions of augmented adrenergic tone. Thus, our findings reveal a mechanistic parsimony showing that HCN4, a channel that contributes to a highly conserved fight or flight heart rate increase, is configured to respond to heat and cAMP. Our findings suggest that heat sensing is core to canonical HCN responses to heat and cyclic nucleotide activation. It will be interesting to learn if HCN channels distributed in brain regions important for systemic thermal control contribute to physiological or disease processes related to increased core body temperature.

## Methods

All experiments involved using animals were carried out in accordance with the guidelines and policies of Institutional Animal Care and Use Committee of Johns Hopkins University.

### SAN cell isolation, culture and infection

Method of isolation of SAN cells from mice was modified and performed according to previously published methods.[26] Mice were anesthetized with avertin (20 μl/g, i.p.) and heparinized (8,000 U/kg, i.p.), and monitored until unresponsive. The heart was excised and placed into ice-cold Tyrode’s solution, consisting of (mM): 140.0 NaCl, 5.0 HEPES, 5.5 Glucose, 5.4 KCl, 1.8 CaCl_2_, and 1.0 MgCl_2_. The pH was adjusted to 7.4 with NaOH. After aortic cannula, the heart was perfused by Ca^2+^-free Tyrode’s solution first, then followed by digestion solution contain collagenase type II (Worthington) and protease type XIV (Sigma). The SAN region, delimited by the crista terminalis, atrial septum and orifice of superior vena cava, was dissected free from the heart and was cut into smaller pieces. SAN tissue pieces were digested in 5 ml of solution containing collagenase type I, elastase (Worthington), and protease type XIV (Sigma) for 20-30 min. The tissue was transferred to 10 ml of Kraft-Bruhe medium containing (mM): 100.0 potassium glutamate, 5.0 HEPES, 20.0 Glucose, 25.0 KCl, 10.0 potassium aspartate, 2.0 MgSO_4_, 10.0 KH_2_PO_4_, 20.0 taurine, 5.0 creatine, 0.5 EGTA, and 1.0 mg/ml BSA, with pH adjusted to 7.2 using KOH. The tissue was agitated using a glass pipette for 10 min. The cells were either kept at room temperature and studied within 7 hours, or used for culture studies. Freshly isolated SAN cells were put into 35 mm tissue culture plates with Media 199 containing Blebbistatin 10 mM (ApexBio B1387-10), 5% FBS, Primocin (Invivogen) and ITS. Fresh medium containing Ad-mCherry-Cre (Vector Biolabs 1773), Ad-HCN4-wt or Ad-HCN4-407/409 was added to the plates at a multiplicity of infection of 100.

### HEK 293 cell culture and transfection

HEK 293FT cells were cultured in DEME (ThermoFisher Scientific, catalog #11995) containing 10% FBS, NEAA (ThermoFisher Scientific, catalog #11140050) and GlutaMax (ThermoFisher Scientific, catalog #35050079). For transfection, cells were seeded between 250,000 to 300,000 cells per 35 mm dish the day before transfection and were then co-transfected with 0.8 μg of an *HCN* plasmid and 0.5 μg of an eGFP plasmid to aid in identifying transfected cells. We used jetPRIME reagent (Polyplus) for transfection.

### Cellular electrophysiology and temperature control

SAN cells or HEK 293FT cells transfected with human HCN4 WT or mutant plasmids were placed in temperature-controlled chamber of around 1 ml volume of Tyrode’s solution at room temperature (around 23±1°C) or warmer temperatures as indicated. Temperature control was performed by TC-344B Dual Channel Automatic Temperature Controller (Warner Instruments, Holliston, MA 01746, USA) used with both a heated chamber and solution in-line heater to rapidly change or maintain a desired temperature. SAN cells with the characteristic morphology (spindle or spider shape) and spontaneous activity were studied. Adenovirus infected SAN cells were identified by mCherry expression. SAN cells were also identified electrophysiologically by typical spontaneous action potentials with slow depolarizing phase 4 and the I*_f_* currents in electrophysiological experiments. HEK 293FT cells with HCN4 plasmids transfection were identified by GFP expression.

Spontaneous action potentials and I*_f_* currents were recorded using the perforated (amphotericin B or β-escin) patch-clamp technique on single SAN cell at various temperatures as indicated in Tyrode’s solution. Temperature was recorded using Digidata 1440A (Molecular Devise, San Jose, CA 95134, USA) and pClamp 11 (Molecular Devise). The patch pipette was filled with (mM): 130.0 potassium aspartate, 10.0 NaCl, 10.0 HEPES, 0.04 CaCl_2_, amphotericin B 240 μg/ml with pH adjusted to 7.2 with KOH. β-escin 50 µM was substituted for amphotericin B to allow dialysis of cAMP or PKI. For recording HCN channels currents in HEK 293FT cells, bath solution composed with (mM): 110.0 NaCl, 0.5 MgCl_2_, 1.8 CaCl_2_, 30.0 KCl, 5.0 HEPES, pH adjusted to 7.4 with NaOH; intracellular solution contains (mM): 130.0 KCl, 10.0 NaCl, 0.5 MgCl_2_, 1.0 EGTA, 5.0 HEPES, pH adjusted to 7.4 with KOH. Data were acquired at 100 kHz using Axopatch 200B (Molecular Devise). When recording I*_f_* currents or HCN4 channels currents, membrane potential was held at −40 mV, the voltage steps were applied for 5 s ranging from −130 mV to −20 mV in 10 mV increments or vice versa. I*_f_* activation analysis was conducted by measuring tail currents [34] and fitting them with Boltzmann equations.

### Ex vivo Langendorff-perfused heart rate measurements

ECG recording from Langendorff-perfused hearts was performed as described [11, 16]. Briefly, excised hearts were rapidly mounted on a Langendorff apparatus (ADInstrments Inc. Colorado Springs, USA) for retrograde aortic perfusion at a constant flow of 4 ml/min with oxygenated (95% O_2_, 5% CO_2_) Krebs-Henseleit solution consisting of (mM): 25.0 NaHCO_3_, 118.0 NaCl, 4.7 KCl, 1.2 MgSO_4_, 1.2 KH_2_PO_4_, 2.5 CaCl_2_, and 11.0 Glucose, with pH equilibrated to 7.4. Each perfused heart was immersed in a water-jacketed bath and the bath temperature was monitored by T-type Implantable Thermocouple Probe (IT-21) (ADInstrments Inc.). ECG measurements from the intact heart were continuously recorded with Isolated Heart MAP Electrode Set (Mouse, ADI) (ADInstrments Inc.). After the heart was allowed to stabilize for at least 20 minutes, the heart rate and bath temperature were recorded by Powerlab (ADInstrments Inc.), bath temperature was controlled or changed by circulating water from bath heater and circulator.

### Cloning and mutagenesis

The plasmid (pENTR223.1-HCN4) containing the human *HCN4* cDNA (HsCD00350637) was ordered from DNASU Plasmid Repository (http://dnasu.org/DNASU/GetCloneDetail.do?cloneid=350637). The *HCN4* cDNA was cloned into the CVM promoter containing expression vector pcDNA6.2-V5-DEST by Gateway cloning (ThermoFisher Scientific). Human *HCN1* (NM_021072.4) and *HCN2* (NM_001194.4) cDNAs were synthesized by TWIST Bioscience and cloned into the pTwist-CMV-BetaGlobin-WPRE-Neo mammalian expression vector. Single and double mutations targeting the candidate residues identified by computational modeling were generated by the Q5 Site-directed mutagenesis kit (New England BioLabs). More complex compound mutations were generated in synthetic gene fragments (TWIST Bioscience), which were then used to replace the corresponding regions in the wildtype plasmids with NEBuilder® HiFi DNA Assembly Cloning Kit (New England BioLabs). All plasmids were fully sequenced. The rabbit M408Q/Y410F (corresponding to M407Q/Y409F on mouse and human HCN4) were generated by the Q5 Site-directed mutagenesis kit.

### HCN4 computational modelling

The transmembrane portion of the open and cAMP-activated structures of HCN4 overlapped using the program MOE (Molecular Operating Environment). The residue substitutions to be studied were modelled in the human structures with the same molecular modelling program. The apolar and polar solvent exposed areas of each atom in the human structures were calculated using a Shrake-Rupley algorithm with a probe radius of 1.4 Å using computational routines provided by Bio.PDB (v 1.79), pandas (v 1.4.1), and Numpy (v 1.22.2) python (v 3.8) modules, summed total for all, main-chain, and side-chain atoms, as well as separated contributions of their polar and apolar areas. The theoretical contributions to the ΔCp of hydration per unit of SASA were calculated using interpolated values at 37°C from table 7 of reference[35]. The linear interpolation used the reported values at 25°C and 50°C. The python Jupyter notebook developed and auxiliary files are available in the GitHub repository (https://github.com/mabianchet/HCN4_SASA).

The comparative model of tardigrade HCN tetramer was obtained using as a template the structure of the human HCN4 (PDBID 6GYN) with the homology modelling routines included in the molecular modelling package MOE, Molecular Operating Environment 2019.01. (2020). The final structure was minimized using an Amber10HT forcefield, also included in MOE. The alignment of Fig. 3e-f was performed with the same program and the figure was prepared with Espript [36]. Structural Figures were prepared with the same program. The model structure is also accessible in (https://github.com/mabianchet/HCN4_SASA).

### M407Q/Y409F knock in mice

*Hcn4*-M407Q/Y409F-knockin mice were generated on a C57BL/6J background using the CRISPR/Cas9 technology at the Johns Hopkins Transgenic Core. A CRISPR guide RNA (Alt-R® CRISPR-Cas9 crRNA, IDT), sequence: 5′-taggtcatgtggaagatctg-3′) was designed to target mouse *Hcn4*. A synthetic ssODN for homology-directed repair was designed to harbor missense mutations for M407Q (ATG→caG), Y409F (TAT→Ttc), and silent mutations encoding I404 (ATC→ATt) and F405 (TTC→TTt) to create an HphI restriction site for genotyping. The sequence of the ssODN was 5’-CAATGAGGTTCACGATGCGTACCACGGCGCTGGCCAGGTCgaAGGTCtgGTGaAAaATCTGTGGAGTCGAGCCATGGCACAGACAAGTAAGCGGTC-3’. Genotyping of founder mice, and subsequent offspring was performed by first PCR amplifying a 385bp fragment from tail DNA with a pair of primers (HCN4QF_Fwd: GTGTTGCCATGGTGCCCTCCAG and HCN4QF_Rev: TGCCACCTGCTGGGATTCAGGA). Then, both Sanger sequencing and HphI digestion of the PCR amplified fragments were used for genotyping. 28 out of 43 F0 mice were targeted by CRISPR, and 3 had correct M407Q/Y409F mutations. Two founder M407Q/Y409F mice were backcrossed with C57BL/6J mice from The Jackson Laboratory, and experiments from both lines produced similar results.

### Statistical analysis

Data are presented as mean±SEM unless otherwise noted. Statistical analysis was performed either with one-way ANOVA or an unpaired or paired Student’s t-test, as appropriate. Analyses were performed with Sigmaplot or Sigmastat (Systat Software, Inc. San Jose, CA 95110 USA), and Graph Pad Prism 8 (GraphPad Software, Inc. San Diego, CA 92108, USA). The null hypothesis was rejected for a P<0.05.

### Movie

Movie 1 was created using the PyMOL program. The intermediate structures between the close and activated form were obtained using The Yale Morphing server (http://www.molmovdb.org/molmovdb/morph/; accessed May 23 2023)[37]

## Movie 1

**Movie 1** depicts the changes in the structure of the human HCN4 during its transitions between a closed and activated state. The orange circle highlights the linker between S4 and S5 that shows significant changes during the hHCN4 activation process. Met 407 and Tyr 409 are part of the linker switch between two distinct environments from near the solvent/lipid interface to be buried in the protein. The red sticks represent these residues.

## Acknowledgement

We appreciate review and suggestions to a draft manuscript by Dr. Francisco Bezanilla of University of Chicago. This work was funded in part by the National Institutes of Health R35HL140034 to (MEA).

## Contributions

Y.W. and M.E.A. conceptualized the study, interpreted the data and wrote the manuscript with the input of co-authors. Y.W., Q.W. and J.G. carried out experiments and analyzed data. Q.W. generated QF mice and most of the experimental reagents. M.B. conducted computational modeling, data interpretation and helped revise the manuscript. A.L. provided HCN4*^fl/fl^* mice and provided input editing the paper. A.M. provided rbHCN4 construct and provided critical input to multiple drafts of the manuscript. O.R.G. generated experimental reagents. E.N.A. generated graphical depictions of HCN4.

---

**Figure S1.**
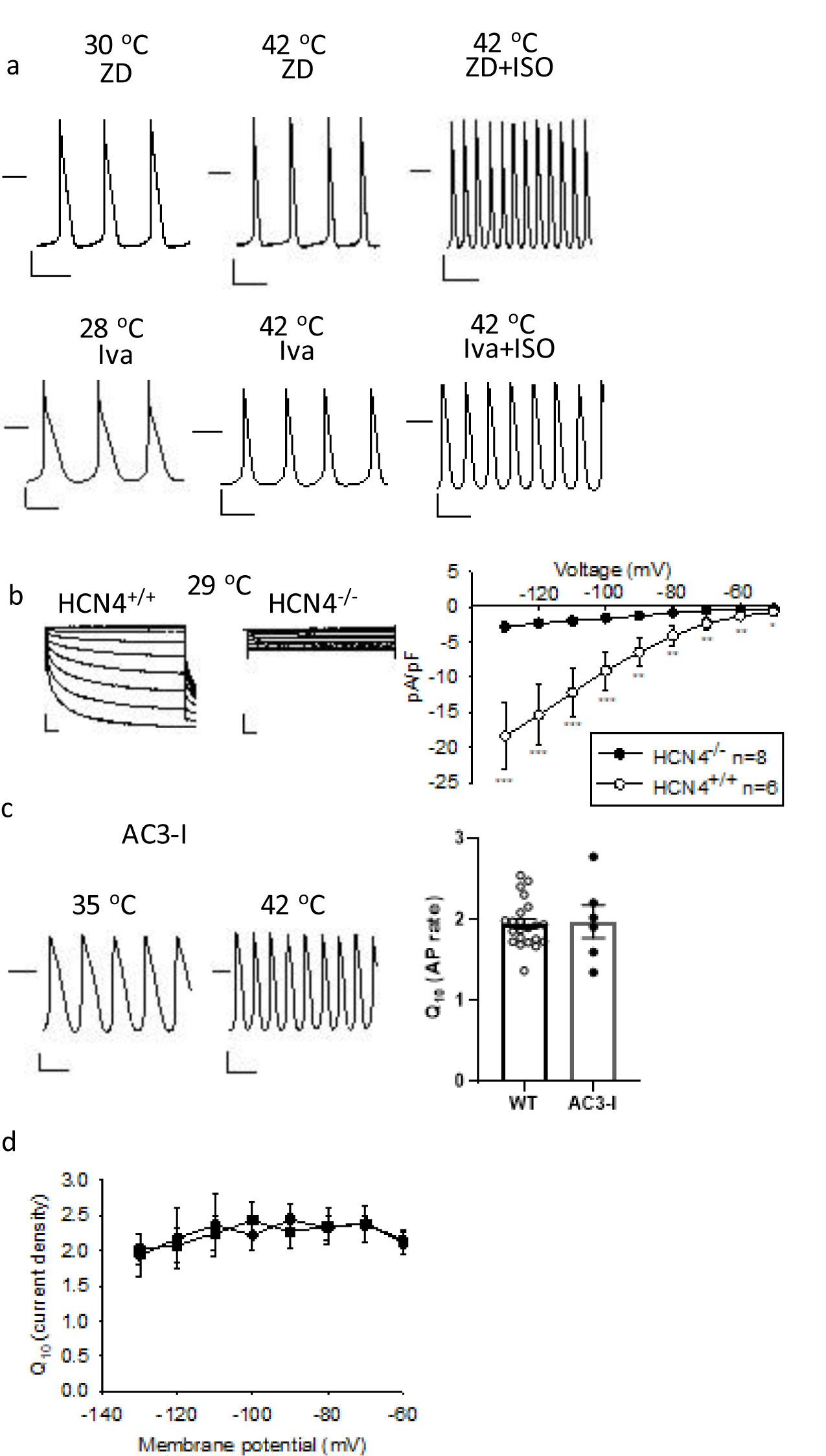
Spontaneous action potentials and HCN4 gene deletion in SAN cells. **a**, Representative spontaneous action potential recordings from isolated mouse SAN cells. ZD7288 (ZD, 4 μM, upper panels) and Ivabradine (Iva, 4 μM, lower panels) reduced temperature dependent, but not ISO (isoproterenol, 1 μM) dependent action potential rate increases. **b,** Representative I*_f_* currents recorded from cultured *Hcn4* floxed (left panel) and cultured *Hcn4* floxed SAN cells co-incubated with Cre recombinase expressing adenovirus (middle panel). Summary data for current-voltage relationships (right panels) from control (left panel, n=6) and *Hcn4* ex se ells (m le p nel, n=). n lys s us n tu ent’s *t* test and Mann-Whitney Rank Sum Test as appropriate. *p<0.05, **p<0.01, ***p<0.001. **c**, Representative action potentials recorded from an SAN cell with transgenic expression of AC3-I, a CaMKII inhibitory peptide at lower (left) and higher (middle) temperatures. Summary Q_10_ data for spontaneous action potential rates are shown on the right. d, Q10 calculated with current densities at testing potential from −60 mV to −130 mV at lower and higher temperatures. Filled circle: If current recorded from single SAN cells, n=11. Filled square: HCN4 channel current recorded from HEK 293 cell transfected with *HCN4*, n=28.

**Figure S2.**
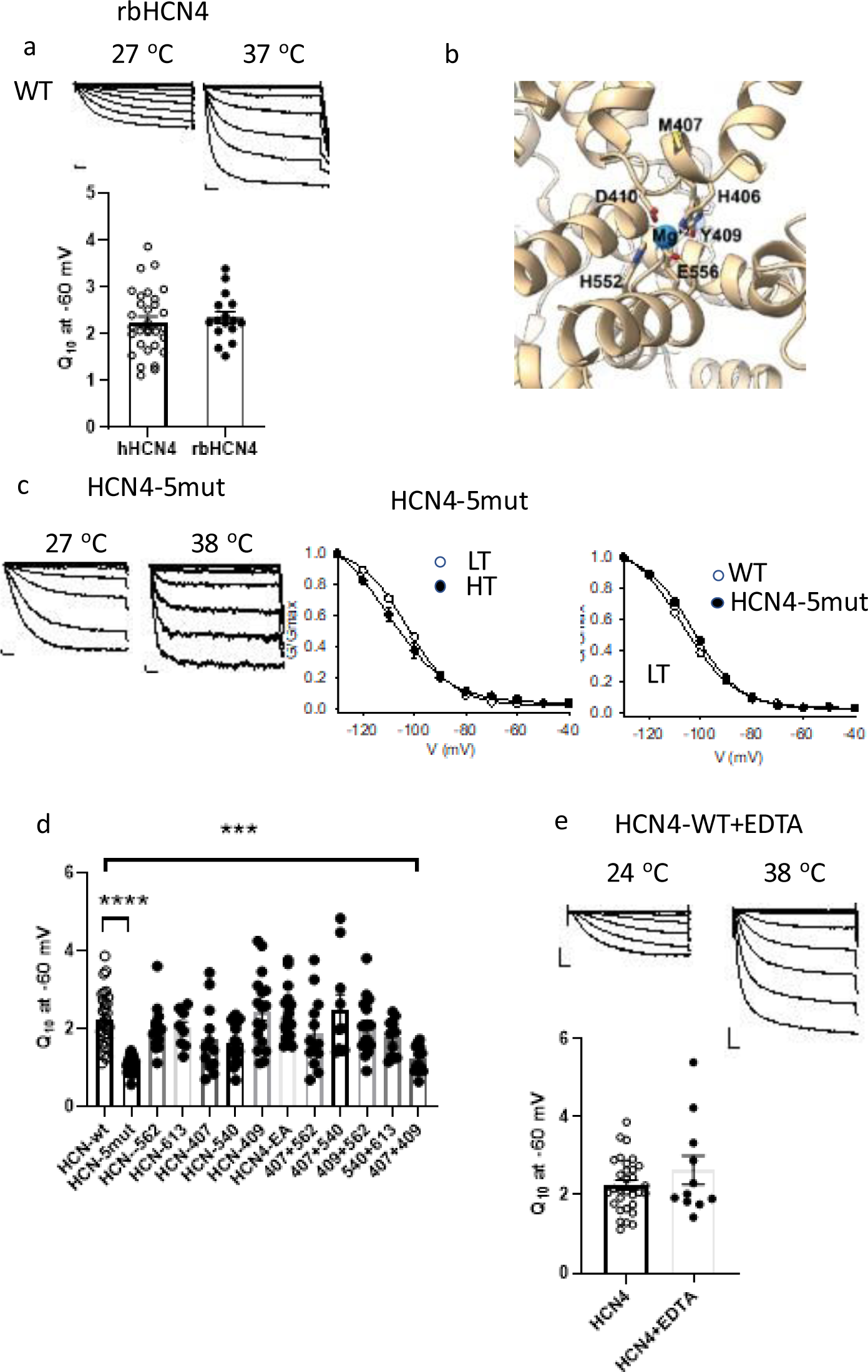
HCN4 heat resistant mutants and structural and functional analysis. **a**, HCN4 channel currents recorded at lower (left) and higher (right) temperature from HEK cell expressing rbHCN4-WT (rabbit WT HCN4) channels. Summary data of Q_10_ at −60 mV from hHCN4 WT compare to rbHCN4 WT (lower panel). **b**, Structural representation of key sites on the S4-S5 intracellular linker and Mg^2+^ coordination sites for the cAMP-activated structure of HCN4 (6GYO), (see Methods). **c,** Representative I*_f_* current traces recorded from a HEK cell expressing HCN4-5mut (see Methods) under low and high temperatures (left two panels). Mean activation Boltzmann fits for I*_f_* under low and high temperatures recorded from HCN4-5mut channels (middle panel, n=45, LT (low temperature range 25-29°C) and n=22, HT (high temperature range 35-41°C) or (right panel) from WT (n=20, LT (low temperature range 25-29°C)) and HCN4-5mut channels (n=45, LT (low temperature range 25-29°C)). **d**, Summary Q_10_ data for I*_f_* recorded at −60 mV from various HCN4 WT and mutant channels (see Methods and Table S2). One way ANOVA test, ***p<0.001, ****p<0.0001. n=8-31/group. **e**, Representative I*_f_* currents recorded with 10 mM EDTA in the pipette solution under low and high temperatures (upper panel). Summary Q_10_ data are shown on the lower panel for normal (n=31) and EDTA (n=11) pipette solutions.

**FigureS3.**
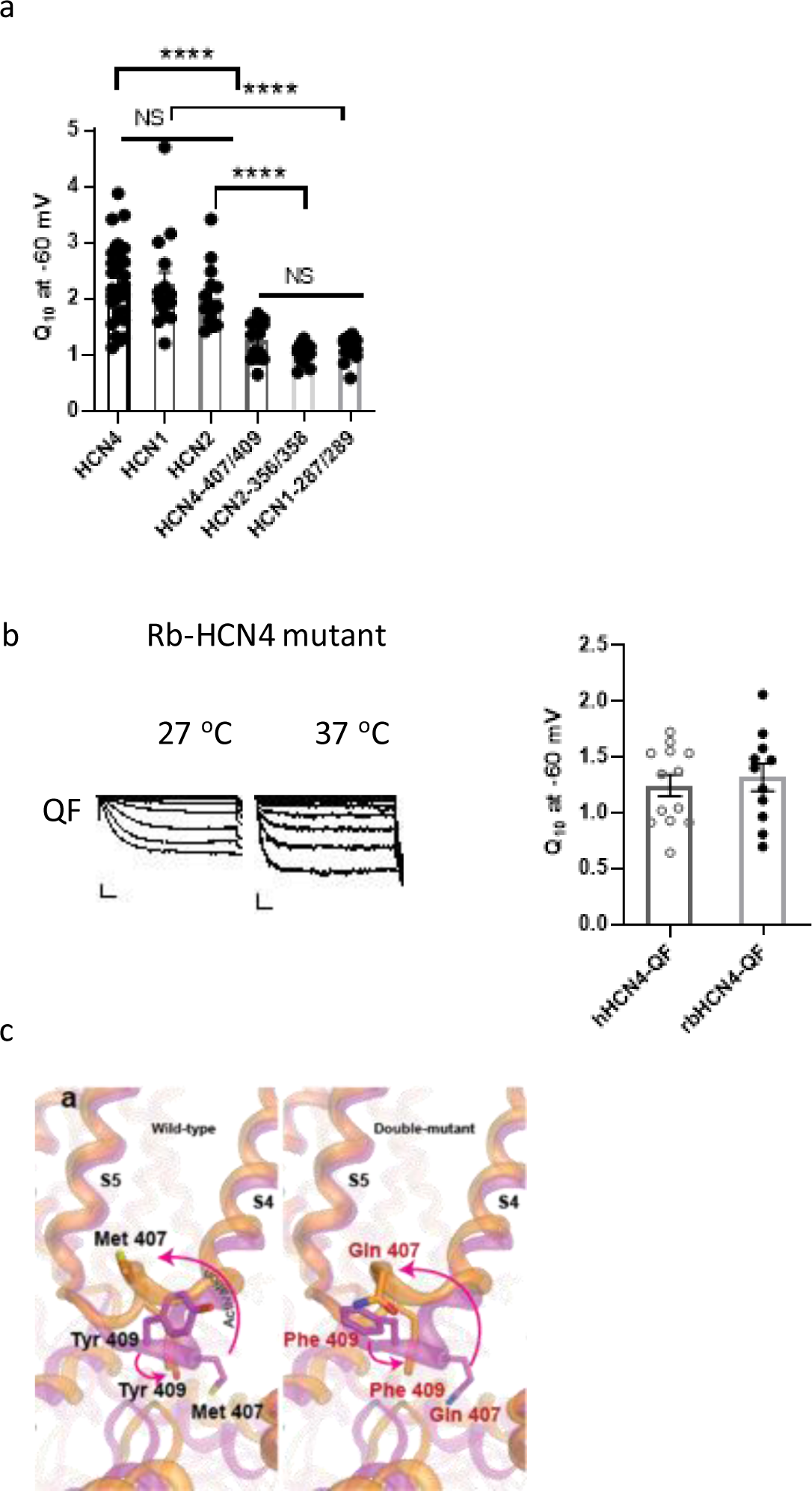
Similar HCN channel family responses to heat. **a**, Q_10_ summary data for I*_f_* recorded at −60 mV from HEK 293 cells expressing WT and heat insensitive mutant HCN1, 2 and 4. One-way ANOVA test, NS, not significant, ****p<0.0001. n=13-31/group. **b,** I*_f_* currents recorded at lower (left) and higher (middle) temperature from HEK cell expressing rabbit rbHCN4-M408Q/Y410F channels. Summary data of Q_10_ at −60 mV from human hHCN4 mutant compare to rbHCN4 mutant on the right. **c**, S4-S5 intracellular linker in the closed and cAMP activated structure. Model of WT Met 407 and Tyr 409 in the closed (magenta) and activated (orange) conformations (left panel). Model of the M407Q and Y409F mutants in closed (magenta) and activated (orange) conformations. The residue rotamers were chosen by a rotational search algorithm by the program MOE. Figures were draw using MOE (see Methods).

**Figure S4.**
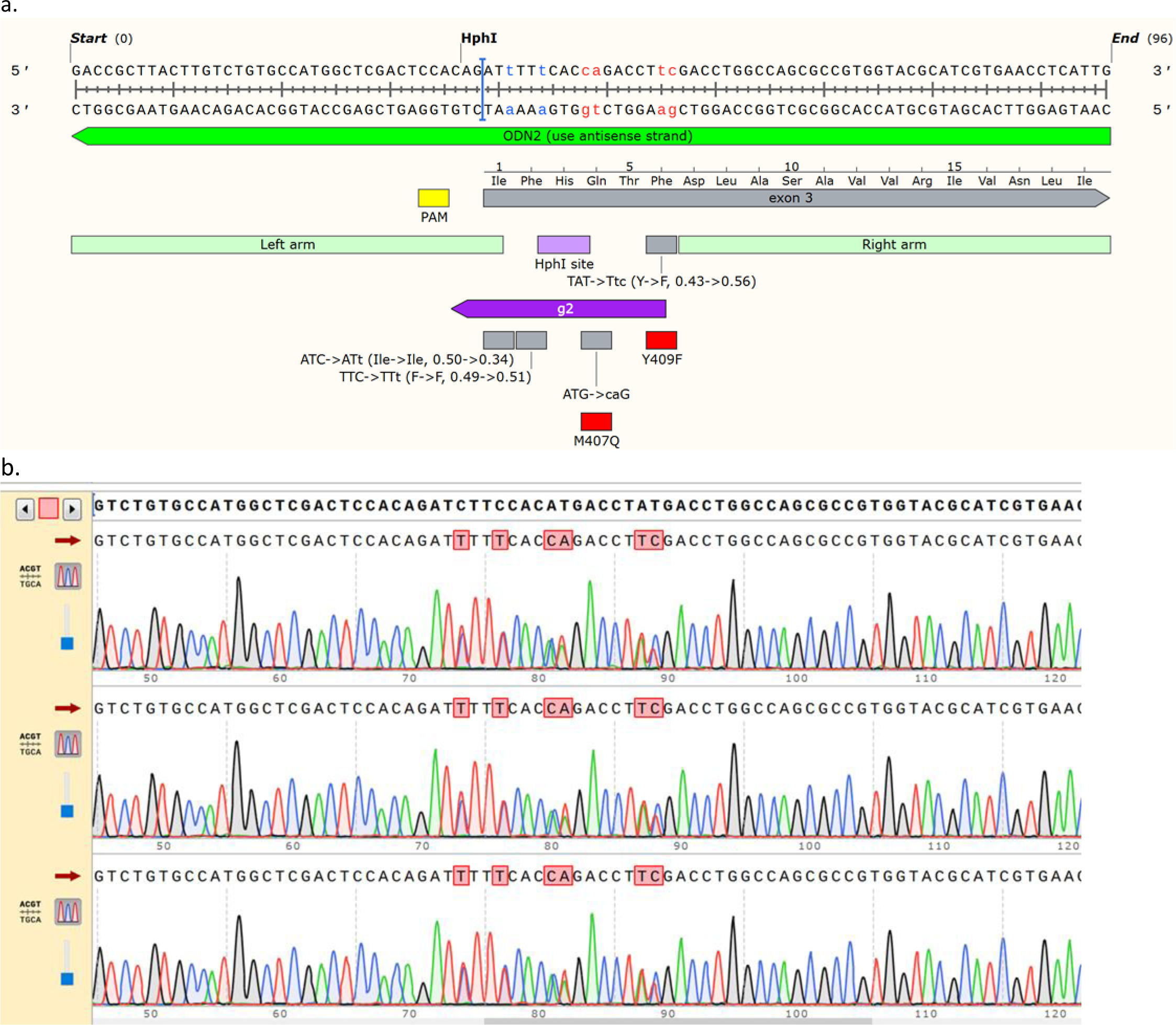
Mouse *Hcn4* CRISPR strategy. **a,** The DNA sequence shows the CRISPR-modified *Hcn4* genomic region. Silent mutations are shown in blue and missense mutations in red. The location of CRISPR guide (g2) binding site, PAM, HphI restriction site, and codons for M407Q and Y409F are indicated. For the silent mutations introduced to the codons for Ile404 and F405, the codon usage frequencies of the original and modified codons are indicated in parentheses. **b,** Sanger sequencing chromatograms of three successfully targeted founder mice.

