## Supplemental File for "HCN channels sense temperature and determine heart rate responses to heat"

**Supplementary Materials**

**Identification of amino acids important for heat sensing**

From a thermodynamic point of view, the free-energy of channel activation by cAMP at pressure and volume constant ($\Delta G^{act.})$is function of specific-heat at constant pressure ($\Delta C_{p}$) of the cAMP-bound and apo (close) states and the temperature, described by equation 1:

$\Delta G^{act.}\left( T \right)= \Delta G^{act.}\left( T_{ref} \right)+\Delta\Delta C_{p}^{act.}\left\{ \left( T-T_{ref} \right)-T\ln\left( \frac{T}{T_{ref}} \right) \right\}$	[eq. 1],

where

$\Delta\Delta C_{p}^{act}=\Delta C_{p}^{cAMP-bound}-\Delta C_{p}^{apo}$	[eq. 2]

The sign of $\Delta\Delta C_{p}^{act.}$ results in a convex (positive) or concave (negative) curvature of $\Delta G^{act.}$ (T), and its magnitude reflects the rate of change with the temperature [1]. A negative $\Delta\Delta C_{p}^{act.}$ will make $\Delta G^{act.}$ more negative (favorable) when the temperature increases (heat-sensitive) and a positive $\Delta\Delta C_{p}^{act.}$ has the opposite effect (cold-sensitive). A lower absolute value of the $\Delta\Delta C_{p}^{act.}$ diminishes the temperature dependence by decreasing the curvature of the $\Delta G^{act.}\left( T \right)$as a function of the temperature.

The Q_10_ value is used to characterize the temperature dependence of the channel activation and is represented using the Arrhenius equations as a function of the energy barrier to overcome during the process ($\Delta E^{\#})$ [2] as:

$Q_{10}=exp\left( \frac{\Delta E^{\#}\left( T \right)}{k_{B}}\left( \frac{10}{T(T+10)} \right) \right)$ [eq. 3],

where k_B_ is the Boltzmann constant, and T the measurement temperature in K. $\Delta E^{\#}$ is the free energy difference between the closed state (initial state) and the highest-energy intermediate (#) of the channel during the reaction path towards the cAMP activated state. At constant pressure and volume, $\Delta E^{\#}$dependence on T is also described by an expression like equation 1:

$\Delta E^{\#}\left( T \right)= \Delta E^{\#}\left( T_{ref} \right)+\Delta C_{p}^{\#}\left\{ \left( T-T_{ref} \right)-T\ln\left( \frac{T}{T_{ref}} \right) \right\}$	[eq. 4],

Channel residues showing significant conformational changes during activation are expected to contribute to establishing the energy barrier that must overcome to become activated ($\Delta E^{\#})$. Importantly, mutation of those residues may significantly affect the channel kinetics and temperature dependence.

To dissect the impact of such mutations on channel Q_10_, we used the ratio between Q_10_ for mutant and wild type, combining equations 3 and 4. This resulted in:

$\frac{Q_{10}^{Mut}(T)}{Q_{10}^{wt}(T)}=\exp\left\{ \left[ \Delta\Delta E^{\#}(T_{ref})+ \Delta{\Delta C}_{p}^{\#mut-wt}\left( \left( T-T_{ref} \right)-T\ln\left( \frac{T}{T_{ref}} \right) \right) \right]\left( \frac{10}{T\left( T+10 \right)} \right) \right\} [\mathrm{eq}.5],$

where $\Delta\Delta E^{\#}$ = $\Delta E^{\# mut}\left( T_{ref} \right)-\Delta E^{\# wt}\left( T_{ref} \right) and \Delta\Delta C_{p}^{\#}=\Delta C_{p}^{\#mut}-\Delta C_{p}^{\#wt}$, choosing an arbitrary $T_{ref}$. Mutations reducing the $\Delta E^{\#}$ with respect to that of the WT results in negative values of $\Delta\Delta E^{\#}$ decreasing the Q_10_ ratio, therefore reducing the mutant Q_10_ Instead, positive values of $\Delta{\Delta C}_{p}^{\#}$ reduces the mutant Q_10_, because of the negative T-explicit term that multiplies $\Delta\Delta C_{p}^{\#}$ in eq. 5.

In proteins, hydration of polar and apolar surfaces largely contribute with opposite signs to the $\Delta G$ of processes in which conformational changes are involved. Protein ${\Delta C}_{p}(T)$ are mostly proportional to the solvent-accessible surface area (SASA) of the protein [3]. Changes in the SASA between apo/closed and cAMP-activated states of the channel contribute to the differential change of both $\Delta E^{\#}$ and $\Delta C_{p}^{\#.}$.

Strategically, by choosing mutations that minimize variations in the residue size, we can limit the structural changes in the mutated protein and neglect the otherwise larger energetic contributions by changing steric interactions when considering the free energy contributions. Mutation changing a residue’s polar or apolar characteristics allows a heuristic estimate of the signs of $\Delta\Delta E^{\#}$ and $\Delta\Delta C_{p}^{\#}$ [1]. The $\Delta\Delta E^{\#}$ sign can be estimated considering the effects on the free energy of hydration of the initial, highest-energy, and final state of the mutation. The structure of the highest-energy intermediate is usually unknown, preventing direct calculation of $\Delta\Delta C_{p}^{\#}$. However, because of the strategy used to choose the mutations, the highest-energy intermediate will maximize the changes by exposure of apolar or sequestering (burying) of polar areas targeted in the mutation. Therefore, the sign of $\Delta\Delta C_{p}^{\#}$ matches the sign of the $\Delta\Delta C_{p}^{mut}=\Delta\Delta C_{p}^{act(mut)}-\Delta\Delta C_{p}^{act(wt)}$ that is calculated from known (or modeled) structures.

**Heat sensing candidate residues**

We calculated the $\Delta\Delta C_{p}^{mut}$ using models of the mutant structures based on the human channel structures (Table S1).

| Table S1. ΔSASA, and $\Delta\Delta C_{p}^{activation}$of residues showing a change in solvent exposure during activation | | | | | | | | | |
| --- | --- | --- | --- | --- | --- | --- | --- | --- | --- |
| Amino acid | **ΔSASA**  **(all) [Å^2^]** | **ΔCp^act^**  **[ J K^-1^ mol^-1^]** | **Mut.** | **ΔΔCp^Mut^**  **[J K^‑1^ mol^-1^]** | **Amino acid** | **ΔSASA**  **(all)**  **[Å^2^]** | **ΔCp^act^**  **[J K^-1^ mol^-1^]** | **Mut.** | **ΔΔCp^Mut^**  **[J K^‑1^ mol^-1^]** |
| Met 407 | 111.9 | 69.5 | M407Q | 11.4 | **Tyr 409** | -141.6 | -141.6 | Y409F | 77.8 |
| Phe 540 | 54.9 | 33.5 | F540Y | 23.2 | **Ile 661** | -99.2 | -202.9 | — | — |
| Lys 562 | -21.7 | 103.3 | K562M | -84.4 | **Arg 668** | -57.8 | 12.0 | — | — |
| Phe 613 | 44.4 | 40.7 | F613Y | -7.2 | **Glu 695** | -83.5 | -22.9 | — | — |

**M407Q.** Met 407 exposes apolar area (81.0 A^2^) during the activation, resulting in an unfavorable hydration enthalpy (Fig. S3c). The exposure this apolar Met contributes positively to the energy of the high-energy intermediate. Mutating this Met to similar size residues, such as Gln, with a smaller apolar area and polar parts, will lower the energy barrier ($\Delta E^{\#})$by increasing the energy of the initial state due to the unfavorable burying of its polar group and decreasing the energy of the intermediate states by the favorable contribution of the solvation of polar parts of the residue. The resulting negative $\Delta\Delta E^{\#}$and positive $\Delta\Delta C_{p}^{\#}$ (Table S1) due to M407Q decreases the Q_10_^mut^.

**Y409F.** Tyr 409 buries 141.6 Å^2^ of SASA between its aromatic ring (76.2 Å^2^ of apolar area) and hydroxyl group (53.2 Å^2^ of polar area) during cAMP activation (Fig. S3c). Y409F mutation eliminates the favorable enthalpy of hydration (ΔΗ < 0) due to the polar area exposure in the apo/closed channel, and the unfavorable enthalpy of hydration of the burying of the tyrosine polar group during the activation. All these result in an increase of the initial state energy, and a potential reduction of the energy of intermediate states, resulting in an overall decrease of ΔE^#^ (and a ΔΔE^#^ < 0). The negative sign ΔΔE^#^ and the positive sign of $\Delta\Delta C_{p}^{\#}$ due to the mutation (Table S1) also reduces the Q_10_^mut^.

***F540Y.*** Phe 540 becomes solvent-exposed (54.9 Å^2^, Table S1) during activation, mostly apolar (ASA_apolar_ = 33.1 Å^2^) with a large positive (unfavorable) contribution to the intermediates free- energy. The F540Y mutation buries in the initial state the tyrosine polar OH group adding a positive contribution to the free-energy of the initial state, and favorable hydration enthalpy during intermediate states to decrease the energy of the highest-energy intermediate. All the above results in negative ΔΔE^#^ and positive $\Delta\Delta C_{p}^{\#}$reducing the Q_10_^mut^. Interestingly, a tyrosine substitution is also observed in tardigrade.

***K562M.*** The channel activation exposes 18.9 Å^2^ of the apolar area and buries 43.8 Å^2^ of the polar area of Lys 562 after activation, both having an unfavorable contribution that may contribute to the high of the energy barrier. Substitution with the more hydrophobic and shorter Met will decrease the unfavorable burying of the Lys polar NH4, resulting in a negative ΔΔE^#^, however, the large negative ΔΔCp^#^ may negate partially or totally reverse the effect of ΔΔE^#^.

***F613Y.*** Similarly, to Phe 540, Phe 613 becomes solvent-exposed during activation (44.4 Å^2^, Table S1). Also, in this case, a substitution with a tyrosine residue will add a polar group to be exposed with a favorable contribution. However, our homology model of the mutant showed that the additional OH group becomes relatively buried, only exposing a small amount of polar area (3.6 Å^2^) resulting in a small negative ΔΔCp^mut-wt^ (Table S1) that may attenuate the effect of ΔΔE^#^ in the Q_10_^mut^.

**Interspecies considerations for modelling**

All HCN4 human (6GYN and 6GYO) and rabbit structures display hydrophobic and hydrophilic interactions of lipids in the S4-linker-S5 region, which may have critical structural functions [4]. Both human and rabbit-activated structures are similar. However, in the human HCN4, the S4-linker-S5 region in the apo/closed state is farther away from the activated state than the apo rabbit structures. The rabbit protein preparations of the Saponaro et al. structural determination[4] used different detergent lipid mixtures to stabilize the rabbit apo cAMP in an open conformation, showing the S4-linker-S5 region conformation closer to the activated state, which is similar in both species.  **This suggest t**hat the linker region presents the largest changes during activation, and we focused our analysis of these changes using human apo and cAMP activated structures.

1.	Chowdhury, S., B.W. Jarecki, and B. Chanda, *A molecular framework for temperature-dependent gating of ion channels.* Cell, 2014. **158**(5): p. 1148-1158.

2.	Sterratt, D.C., *Q10: The Effect of Temperature on Ion Channel Kinetics*, in *Encyclopedia of Computational Neuroscience*, D. Jaeger and R. Jung, Editors. 2013, Springer New York: New York, NY. p. 1-3.

3.	Makhatadze, G.I. and P.L. Privalov, *Heat capacity of proteins: I. Partial molar heat capacity of individual amino acid residues in aqueous solution: Hydration effect.* Journal of Molecular Biology, 1990. **213**(2): p. 375-384.

4.	Saponaro, A., et al., *Gating movements and ion permeation in HCN4 pacemaker channels.* Mol Cell, 2021. **81**(14): p. 2929-2943 e6.
