## Supplemental Table for "HCN channels sense temperature and determine heart rate responses to heat"

**Table S1.** ΔSASA, and ΔΔCp^activation^ of residues showing a change in solvent exposure during activation.

| Amino acid | ΔSASA (all) Å^2^ | ΔCp^act^ J K^-1^ mol^-1^ | Mut. | ΔΔCp^Mut^ J K^-1^mol^-1^ |
| --- | --- | --- | --- | --- |
| Met 407 | 111.9 | 69.5 | M407Q | 11.4 |
| Phe 540 | 54.9 | 33.5 | F540Y | 23.2 |
| Lys 562 | -21.7 | 103.3 | K562M | -84.4 |
| Phe 613 | 44.4 | 40.7 | F613Y | -7.2 |
| Tyr 409 | -141.6 | -141.6 | Y409F | 77.8 |
| Ile 661 | -99.2 | -202.9 | — | — |
| Arg 668 | -57.8 | 12.0 | — | — |
| Glu 695 | -83.5 | -22.9 | — | — |

**Table S2. Activation curve parameters**

|  | V1/2 (mV) ±SEM  control | K (mV) ±SEM  control | n | V1/2 (mV) ±SEM  +cAMP | K (mV) ±SEM  +cAMP | n | cAMP-induced  V1/2 shift±SEM | Temp-induced  V1/2 shift±SEM |
| --- | --- | --- | --- | --- | --- | --- | --- | --- |
| HCN4 wt (LT) | -107.3±0.7 | 10.4±0.7 | 20 | -96.1±1.2 | 14.1±1.3 | 14 | 11.2±1.4^***^ |  |
| HCN4 wt (HT) | -96.0±1.4^###^ | 10.5±0.8 | 21 | -91.1±2.8 | 11.1±1.2 | 13 | 4.9±3.1 | 11.3±1.6^***^ |
| HCN4 M407Q-Y409F (LT) | -105.3±1.2 | 8.9±0.4 | 27 | -104.6±1.6 | 9.1±1.3 | 13 | 0.7±2.0 |  |
| HCN4 M407Q-Y409F (HT) | -108±1.5 | 10.7±0.8 | 13 | -103.2±2.3 | 8.4±0.9 | 10 | 4.8±2.7 | -2.7±1.9 |
| HCN4 M407Q-Y409F-F540Y-K562M-F613Y (LT) | -102.7±1 | 8.3±0.3 | 45 |  |  |  |  |  |
| HCN4 M407Q-Y409F-F540Y-K562M-F613Y (HT) | -108.2±2 | 9.9±0.7 | 22 |  |  |  |  | -5.5±2.2^**^ |
| HCN4 R669E-T670A (LT) | -105.2±1.5 | 11.5±0.8 | 22 | -107.3±1.9 | 11.1±0.8 | 11 | -2.1±2.4 |  |
| HCN4 R669E-T670A (HT) | -97.9±1.6^###^ | 10.7±1.0 | 18 | -102.0±3.7 | 9.0±1.3 | 7 | -4.1±4.0 | 7.3±2.2^**^ |
| HCN4 Y527F-R669E-T670A (LT) | -95.9±1.5^###^ | 11.4±0.6 | 12 | -98.6±1.2 | 11.1±0.6 | 13 | -2.7±1.9 |  |
| HCN4 Y527F-R669E-T670A (HT) | -96.3±1.9^###^ | 9.9±1 | 9 | -97.2±2.8 | 12.2±0.9 | 5 | -0.9±3.4 | -0.4±2.4 |
| HCN4 wt with EDTA (LT) | -101.4±1.6 | 13.7±0.7 | 13 |  |  |  |  |  |
| HCN4 wt with EDTA (HT) | -95±1.4^#^ | 12±0.8 | 12 |  |  |  |  | 6.4±2.1^**^ |
| HCN1 wt (LT) | -84.9±1.6 | 9.1±0.6 | 16 | -84.3±4.6 | 13.7±3.9 | 3 | 0.6±4.9 |  |
| HCN1 wt (HT) | -91.2±1.8^#^ | 11.1±0.9 | 15 | -83.3±3.9 | 9.8±1.4 | 3 | 7.9±4.3 | -6.3±2.4^*^ |
| HCN1 M287Q-Y289F (LT) | -96.7±2.7^###^ | 12.4±0.8 | 9 |  |  |  |  |  |
| HCN1 M287Q-Y289F (HT) | -100.0±2.9^###^ | 12.0±0.6 | 6 |  |  |  |  | -3.3±4.0 |
| HCN2 wt (LT) | -97.4±1.5 | 10.0±0.6 | 32 | -90.0±1.9 | 9.6±0.7 | 16 | 7.4±2.4^**^ |  |
| HCN2 wt (HT) | -87.2±2.5^###^ | 9.6±0.9 | 18 | -81.0±2.8 | 10.2±0.9 | 10 | 6.2±3.7 | 10.2±2.9^***^ |
| HCN2 M356Q-Y358F (LT) | -107.9±1.3^###^ | 9.9±0.9 | 19 | -107.3±0.8 | 9.1±0.7 | 16 | 0.6±1.5 |  |
| HCN2 M356Q-Y358F (HT) | -107.0±2.5^###^ | 14.0±1.2 | 12 | -108.0±3.5 | 13.0±1.5 | 8 | -1.0±4.3 | 0.9±2.8 |
| HCN2 R618E-T619A (LT) | -102.8±1.2^##^ | 8.7±0.4 | 25 | -103.4±0.9 | 8.8±0.4 | 22 | -0.6±1.5 |  |
| HCN2 R618E-T619A (HT) | -104.7±1.0^##^ | 9.9±0.8 | 16 | -101.7±1.9 | 9.4±0.7 | 15 | 3±2.1 | -1.9±1.6 |

Half-activation voltage (V1/2) and inverse slope factor (k) obtained by fitting data to a Boltzmann function (see Methods) in absence or presence of cAMP (1 mM); LT (Low temperature, 22-29 ^o^C) and HT (High temperature, 35-43 ^o^C); n=number of cells tested in each condition.

###p<0.001, ##p<0.01, #p<0.05 by One-way ANOVA with Multiple Comparisons versus Control Group (Holm-Sidak method) compared to wt HCN4, HCN1 or HCN2 at LT; ***p<0.001, **p<0.01, *p<0.05 by Student’s *t*-test compared to control condition (cAMP compare with without cAMP or HT condition compare with LT condition).
